# A novel RofA-family transcriptional regulator, GadR, controls the development of acid resistance in *Listeria monocytogenes*

**DOI:** 10.1101/2023.06.22.546042

**Authors:** Jialun Wu, Olivia McAuliffe, Conor P O’Byrne

## Abstract

Stomach acid provides a significant innate barrier to the entry of the food-borne pathogen *Listeria monocytogenes* into the human gastrointestinal tract. A key determinant of acid resistance in this bacterium is the conserved glutamate decarboxylase system, GadD2 (encoded by the *gadT2D2* operon), which helps to maintain the intracellular pH during exposure to gastric acid. In this study, we identified a premature stop codon in a gene located immediately downstream of the *gadT2D2* operon that was highly linked to an acid sensitive phenotype. When this open reading frame was restored through homologous recombination an acid resistant phenotype resulted. Through a series of genetic, transcriptomic and survival experiments we established that this gene, which we designated *gadR*, encodes a transcriptional regulator of the *gadT2D2* operon. GadR belongs to the RofA family of regulators, primarily found in the streptococci, where they are involved in regulating virulence. The data further showed that *gadR* plays a critical role in the development of acid resistance in response to mild acid exposure, a response that is known as the adaptive acid tolerance response (ATR). A deletion analysis of the *gadT2D2* promoter region identified two 18bp palindromic sequences that are required for the GadR-mediated induction of *gadT2D2*, suggesting that they act as binding sites for GadR. Overall, this study uncovers a new RofA-like regulator of acid resistance in *L. monocytogenes* that plays a significant role in both growth phase-dependent and ATR mediated acid resistance and accounts for previously observed strain-to-strain differences in survival at low pH. The findings have important implications for understanding the behavior of *L. monocytogenes* in acidic environments and identify a potential target for improved control of this important pathogen.

**Author summary:** The ability to survive the acidic conditions found in the stomach is a key trait enabling the food-borne pathogen *Listeria monocytogenes* to gain access to mammalian gastrointestinal tract, where it can initiate an infection. Little is currently known about how acid resistance is regulated in this pathogen and why this trait is highly variable between strains. Here we used genomic sequences from a collection of *L. monocytogenes* strains with known differences in acid survival to identify a novel transcriptional regulator controlling acid resistance, which we call GadR. The regulator belongs to a family of regulators previously found only in a small group of bacterial pathogens including the streptococci, where they are involved in regulating virulence properties. We show that GadR is the dominant regulator of acid resistance in *L. monocytogenes* and that variability in its gene sequence accounts for previously observed differences between strains in this trait. Together these findings significantly advance our understanding of how this important pathogen copes with acid stress and suggests a potential molecular target to better control it in the human food-chain.

## Introduction

The bacterium *Listeria monocytogenes* is a well-studied member of the *Bacillota* phylum (formerly the *Firmicutes*) due to its impact on food safety and its behavior in the host as a facultative intracellular pathogen. Ingestion of food contaminated with this bacterium is associated with a risk of infection, termed listeriosis, especially in immunocompromised individuals, where the mortality rate can be as high as 30% [1–3]. The organism is particularly problematic for producers of ready-to-eat foods as it has an ability to grow at refrigeration temperatures and in some foods preserved at low pH and/or low water activity [4–6]. Its robust response to acid stress also increases the risk that the pathogen can survive transit of the acidic conditions in the stomach and thereafter invade the epithelium of ileum [7,8], which is the first stage of pathogenesis during an infection [3]. While much has been learned about the mechanisms that contribute to the acid stress response in this pathogen [4,9–11], the regulatory mechanisms that control the expression of these systems are much less well understood.

Almost three decades ago, an adaptive response to acid was described in *L. monocytogenes*, whereby a mild acid stimulus (pH 4.0-6.0) triggers the development of high levels of acid resistance to normally lethal acid conditions (pH 3.0) [12,13]. This response, which was named the Adaptive acid Tolerance Response (ATR), requires *de novo* protein synthesis but the mechanisms underpinning its regulation have never been properly elucidated. Interestingly, entry to stationary phase was also found to promote the acid resistance in *L. monocytogenes* independently of pH [12]. The discovery that Sigma B (SigB), an alternative sigma factor that controls the general stress response in *L. monocytogenes*, plays an important role in acid resistance [14–17] initially suggested that it might play a role in regulating stationary phase acid resistance and the ATR. SigB is activated in stationary phase, it responds to mild acid stress, and it plays a role in transcribing key components involved in protection against acid stress, including the glutamate decarboxylase and arginine deiminase pH homeostasis systems [17–19]. Furthermore, mutants lacking SigB have an acid-sensitive phenotype in stationary phase and a reduced ATR [14,16,19]. However, *sigB* mutants are still capable of developing an increase in acid resistance in stationary phase and inducing a significant ATR [16,19], suggesting that other regulatory factors must be involved in regulating the acid stress response of this pathogen.

Two glutamate decarboxylases (GadD2 and GadD3) are present in most strains of *L. monocytogenes*, and a third (GadD1) is present in some lineage II strains (Fig. 1*A*). The corresponding genes for these decarboxylases are encoded at three distinct genetic loci [20–23]. They contribute to acid stress protection by helping to maintain the intracellular pH, because the decarboxylation reaction consumes protons [24–26]. Two of these systems are genetically coupled with Glu/GABA antiporters (GadD1T1 and GadT2D2) that allow uptake of glutamate (Glu) in exchange for the product of the decarboxylation reaction, gamma aminobutyrate (GABA). GadD3 is not co-expressed with an antiporter and appears to use intracellular glutamate independently of a Glu/GABA exchange mechanism [27,28]. GadD1T1 is reported to contribute to growth under acidic conditions [21] whereas GadT2D2 is the dominant system with respect to acid resistance [23,29]. Little is known about the transcriptional regulation of these two systems, although the transcription of *gadD3* is under SigB control [30,31]. SigB also controls the expression of succinate semialdehyde dehydrogenase (Lmo0913), which is required in GABA-shunt pathway that generates succinate and Glu from GABA and α-ketoglutarate [15,32]. There are significant strain-to-strain differences in the behaviors of the GAD systems in *L. monocytogenes* [23,28]. In particular, the well-studied lab strains EGD-e and 10403S, both of which belong to lineage II serotype 1/2a, produce and secrete different amounts of GABA in response to acidification. Specifically, 10403S secretes GABA in response to acidification whereas EGD-e does not, although *gadD1T1*, *gadT2D2*, and *gadD3* are conserved in these two strains [23]. Furthermore, deletion of each of three GAD systems from these two strains produces different phenotypes with respect to acid resistance; notably deletion of *gadD2* produces an acid sensitive phenotype in 10403S but has no effect on EGD-e [23]. It is striking that the well-studied lab strain EGD-e is especially sensitive to acid, the genetic basis for which has remained unclear [23,28,29].

**Fig 1.**
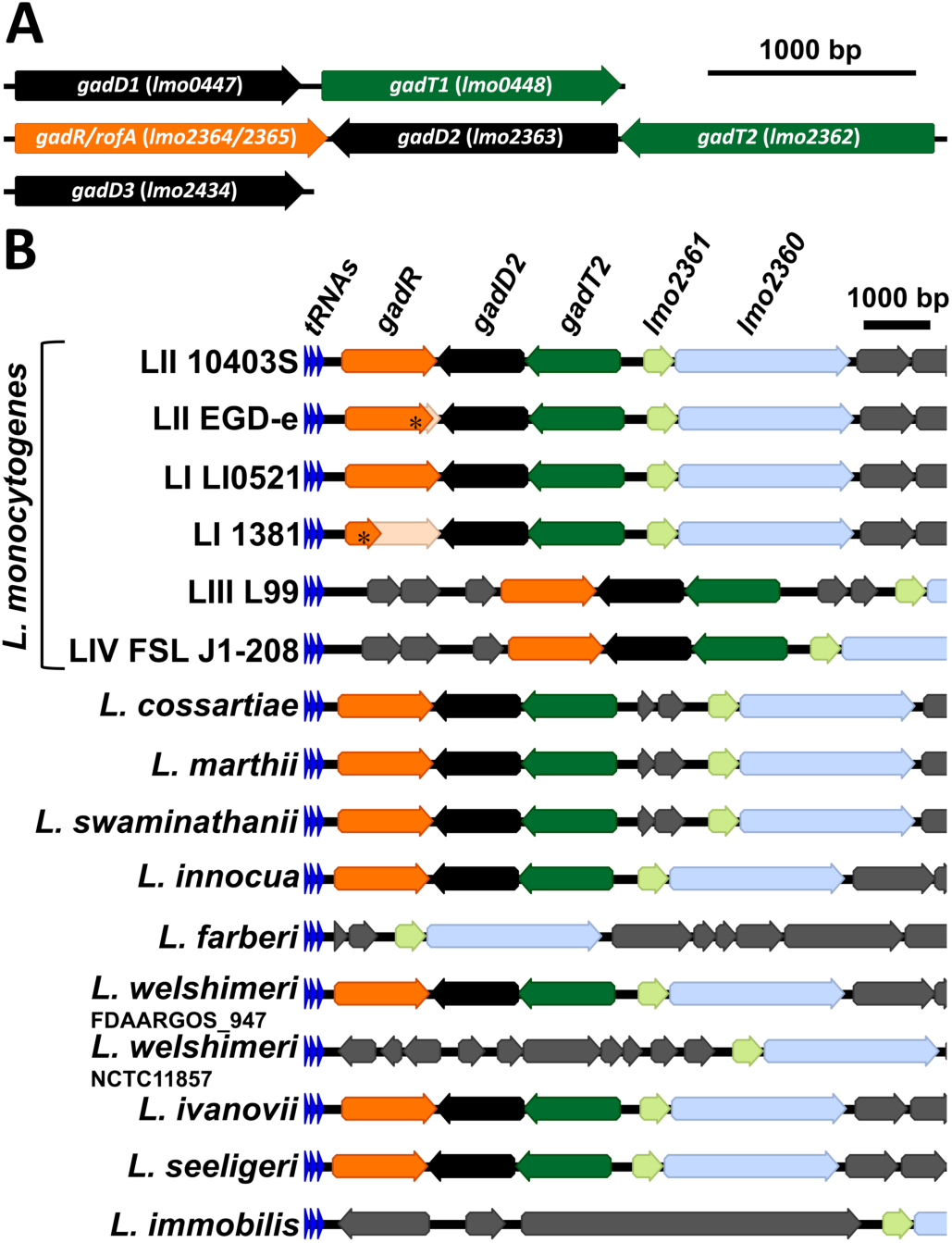
The *gadT2D2-gadR* gene cluster is highly conserved in the *Listeria senso stricto* clade. (A) Operon structures of *gadD1T1, gadT2D2-gadR*, and *gadD3* are depicted, using sequences were extracted from strain 10403S. (B) Gene arrangements downstream of the conserved *tRNA-Gly* are depicted for all species from *Listeria senso stricto* clade species reported by Carlin et al. [36]. *L. monocytogenes* lineage I-IV are abbreviated as LI-LIV. The PMSCs in *gadR* are highlighted with *, the untranslated regions of *gadR* are faded.

We have recently described the phylogenetic and phenotypic characterization of a collection of 168 lab, food, environmental and clinical isolates of *L. monocytogenes* with respect to their acid tolerance (growth at pH 4.9, to not confuse with ATR), acid resistance (survival at pH 2.3) and salt tolerance (growth in NaCl 0.8 M to 1.5 M) [33]. This collection is both phylogenetically and phenotypically diverse, with wide variabilities in acid tolerance and acid resistance across individual strains. Some phenotypic outliers were identified as harboring lesions in the *sigB* operon, which explains compromised general stress response and acid sensitive phenotypes [33]. One strain identified in that study, a lineage I strain designated 1381, displayed reduced acid tolerance (no growth at pH 4.9) and reduced acid resistance (poor survival at pH 2.3) although there was no evidence of mutations affecting the SigB system [33]. We have recently shown that the growth defect in strain 1381 at low pH, but not the poor survival phenotype (at pH 2.3), is caused by a mutation in the *mntH* gene, which encodes a manganese transporter belonging to the NRAMP family [34]. However, this mutation does not account for the decreased acid resistance in this strain [34]. Thus, the genotype of this strain could help provide new insights into the mechanisms underpinning acid resistance in *L. monocytogenes*.

In the present study, we used comparative genomics to help further elucidate the genetic basis for the reduced acid resistance observed in this strain. Our findings reveal the presence of a hitherto unknown transcriptional regulator, GadR, that plays a crucial role in the regulation of acid resistance by modulating the expression of the GadT2D2 in this pathogen. The presence of a mutation within the *gadR* gene is shown to solely account for the differences in acid resistance between the two well-studied lab strains, EGD-e and 10403S. Overall, this study identifies a key regulatory component of adaptive acid resistance in *L. monocytogenes* and raises the possibility that acid resistance, and thus virulence, might be controlled through a strategy that specifically targets this regulator.

## Results

### Identification of a RofA-like regulator that positively influences acid resistance

The acid sensitive and acid intolerant CC2 strain 1381 is closely related to the chromosomally sequenced reference strain LI0521 [34], facilitating an investigation into the genetic basis of these unusual phenotypes. There are 4 genes intact in strain LI0521 that are truncated by the presence of premature stop codon (PMSC) in strain 1381, including *mntH* [34]. Among the other three is a putative RofA-like transcriptional regulator, located downstream of and oriented convergently with the acid resistance operon *gadT2D2* (Fig 1). Notably, this gene was also truncated in the widely studied lab strain EGD-e (L374*) and thus mis-annotated as two open reading frames (ORF), *lmo2365 and lmo2364* in that strain [35]. Bioinformatic analysis revealed that this *gadT2D2-rofA* gene cluster is conserved in most species from *Listeria sensu stricto* clade (Fig 1B) [36] but absent from the *Listeria sensu lato* clade (data not shown). These observations suggested that this RofA-like transcriptional regulator might be the cognate regulator of the *gadT2D2* operon and therefore, we designated the full ORF as *gadR*. Interestingly, in addition to strains 1381 and EGD-e, CC7 strain 1147 and all five CC18 strains from the previously characterized strain collection (n = 168) are predicted to be *gadR*^-^ based on the presence of PMSCs within the ORF (data not shown). All these *gadR*^-^ strains survived poorly at pH 2.3 [33], suggesting a possible role of GadR in acid resistance.

To test whether GadR influences acid resistance and *gadT2* transcription, full length *gadR* was cloned from CC2 strain 1380 and introduced into strain 1381 using an IPTG-inducible integrative expression vector pIMK3, generating construct pIMK3::*gadR* [37] (Table 1). The acid resistance of strain 1381 pIMK3::*gadR* during stationary phase was enhanced in an IPTG-dependent fashion compared to strain 1381 (Fig 2A). The transcription levels of *gadT2* were positively correlated with the presence of a functional copy of *gadR* (from pIMK3::*gadR*) (Fig 2B and 2C). The pIMK3 vector did not affect acid resistance or *gadT2* transcription (Fig 2A and 2B). These results demonstrated that GadR positively influences both acid resistance and *gadT2* transcription. When the pIMK3::*gadR* plasmid was transformed into the *mntH*^+^ strain 1381R1 [34] (Table 1), the positive effects on both acid resistance and *gadT2* transcription were very similar to that observed in the *mntH*^-^ parental strain (Fig 2A and 2B), suggesting that the effect of GadR on acid resistance is independent of MntH.

**Fig 2.**
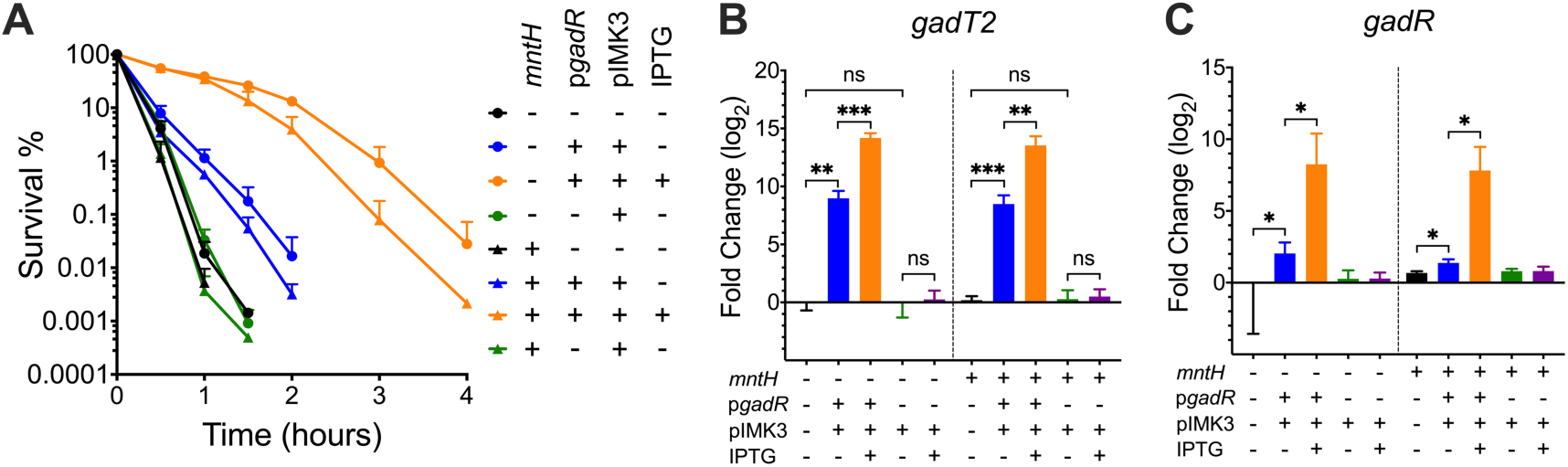
GadR positively influences acid resistance and *gadT2* transcription, independently of the manganese transporter MntH. (A) Stationary phase cultures of strains 1381 (*mntH*^-^) and 1381R1 (*mntH*^+^) either with or without *gadR* complementation (p*gadR*) and 1 mM IPTG induction were challenged at pH 2.3 and survival recorded over 4h. Transcription of *gadT2* (B) and *gadR* (C) was measured for these strains after overnight incubation for 18 h. Transcript levels relative to the *gadR*^-^ parent strain 1381 are presented. Statistically significant differences were determined by paired *t* test (two-tailed) (ns, not significant; * *p* < 0.05; **, *p* < 0.01; ***, *p* < 0.001).

**Table 1.**
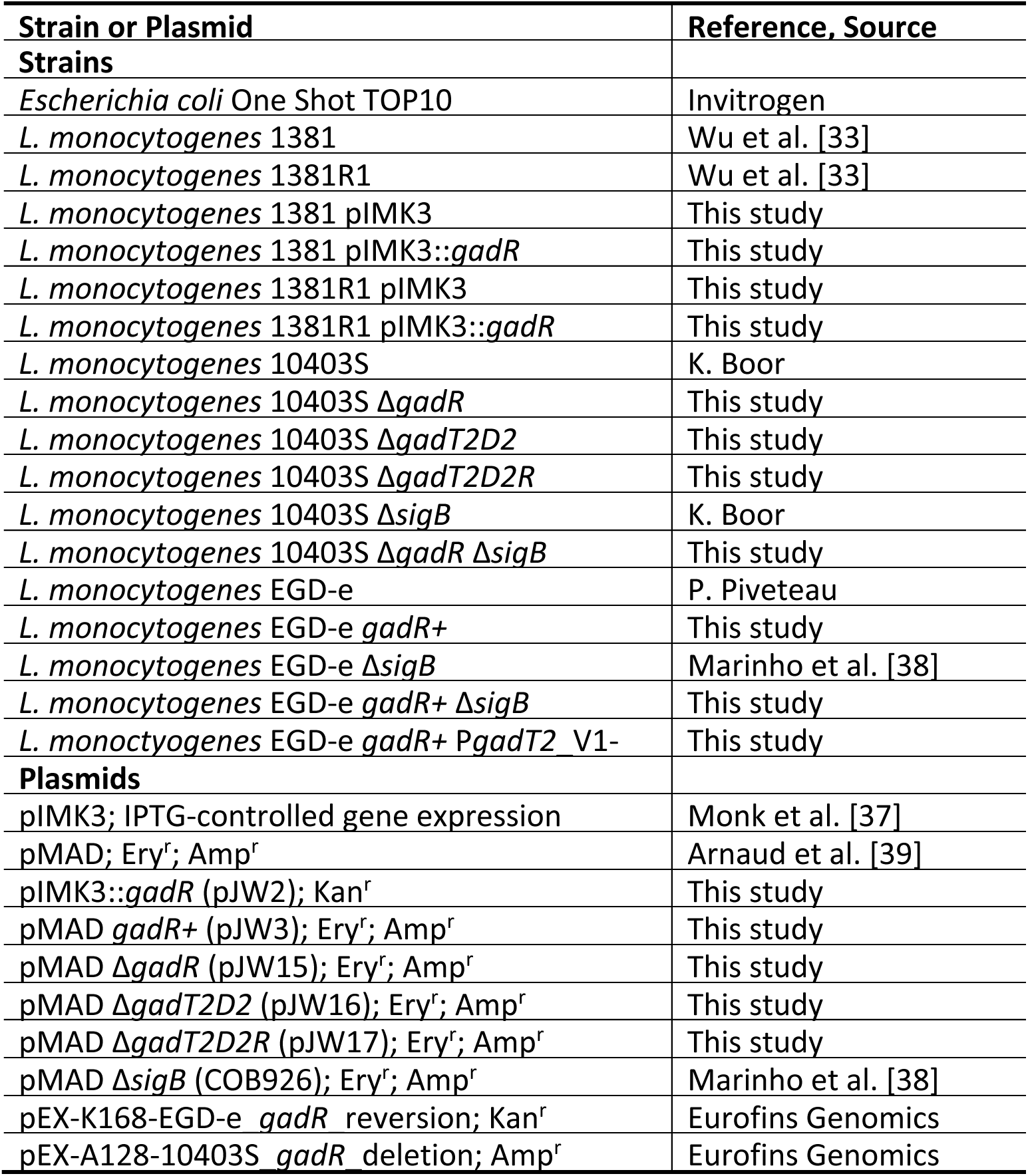
Strains and plasmids used in this study.

### GadR controls acid resistance by activating GadT2D2 expression independently of SigB

To test whether altered *gadT2D2* transcription is exclusively responsible for GadR mediated acid resistance, three mutant strains were constructed in a genetic background where the *gadR* was intact (lab strain 10403S); they carried deletions either in the *gadR* gene (Δ*gadR),* the *gadT2D2* locus (Δ*gadT2D2)* or both (Δ*gadT2D2R*) (Table 1). The ability of these strains to survive at pH 2.3 was compared following growth to stationary phase. All three mutants exhibited a very similar decrease in acid resistance compared to the parental strain 10403S (Fig 3A), suggesting that the loss of GadT2D2 expression is solely responsible for the GadR-mediated acid resistance. The GadR-mediated acid resistance in strain EGD-e (*gadR*^-^) was restored by repairing the point mutation in *gadR* (*374L) in the chromosome by homologous recombination (Fig 3A and 3B). These observations demonstrated a prominent role for GadR in acid resistance and indicated that the *gadR* frameshift mutation in EGD-e explains the intrinsic acid sensitivity of this reference strain. To examine the relative contributions of GadR and the general stress response regulator SigB to acid resistance, Δ*sigB* was introduced into strains 10430S and EGD-e with or without *gadR* (Table 1). And the acid resistance of strains lacking either or both of *gadR* and *sigB* was compared to the strains that have functional SigB and GadR in both genetic backgrounds (strains EGD-e and 10403S). The strains EGD-e *gadR*^-^ Δ*sigB* and 10403S Δ*gadR* Δ*sigB* were extremely acid sensitive and therefore, the cultures were grown to stationary phase and exposed to pH 2.4 (rather than pH 2.3). In both genetic backgrounds, GadR made a more significant contribution than SigB to acid resistance, while the absence of both regulators resulted in extreme acid sensitivity (Fig 3B). These results suggested that the combined effects of GadR and SigB accounts for most, if not all, of the acid resistance of *L. monocytogenes* at pH 2.4 in stationary phase. To elucidate possible crosstalk between GadR and SigB, the transcription levels of *gadT2*, *gadR*, and the SigB dependent *gadD3* during stationary phase were measured for these strains (Fig 3C-3E). The results suggested that SigB has limited, if any, influence on GadR-mediated *gadT2* transcription. *gadR* transcription was generally stable across these strains suggesting that it is controlled neither by GadR nor by SigB (Fig 3D). The transcription of *gadD3* was confirmed to be SigB-dependent and was unaffected by *gadR* deletion (Fig 3E), suggesting that GadR does not influence SigB activity and does not control *gadD3* transcription. To establish if the GadR-mediated *gadT2* transcription is associated with altered Glu/GABA antiporter activity, the ability to export GABA into the medium was examined for these strains during stationary phase. The results showed that significant extracellular GABA was only detectable in the presence of both *gadR* and *gadT2D2* (Fig 3F). Taken together, these results demonstrated that GadR promotes acid resistance by activating the expression of the GadT2D2 independently of SigB.

**Fig 3.**
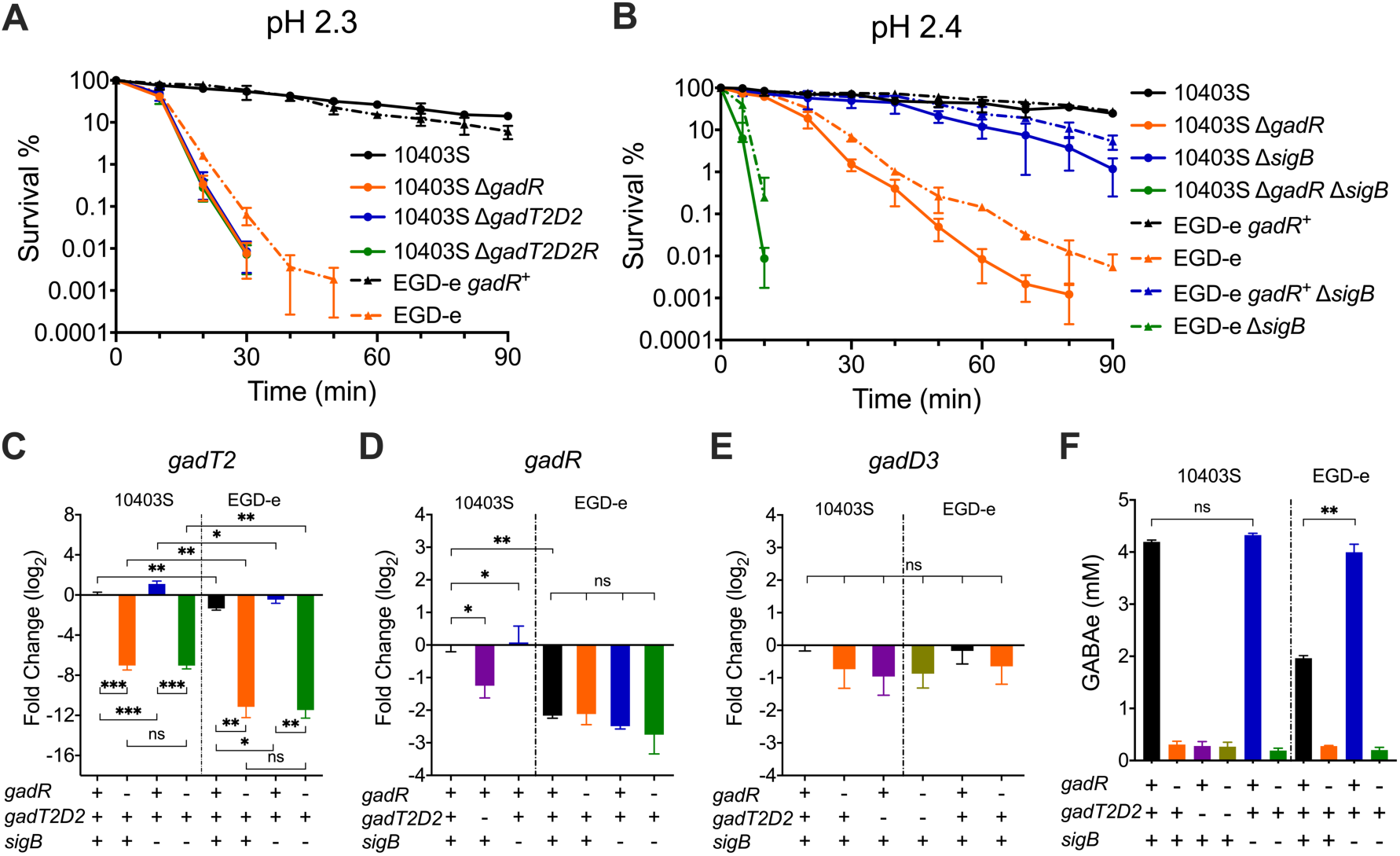
GadR modulates acid resistance by activating GadT2D2 expression independently of SigB. Cultures of *L. monocytogenes* WT or mutant strains were grown to stationary phase and challenged at pH 2.3 (A) or pH 2.4 (B). EGD-e carries a frameshift in the *gadR* gene whereas the ORF was restored to full-length by homologous recombination in the strain EGD-e *gadR*^+^. Transcription of *gadT2* (C), *gadR* (D), and *gadD3* (E) in these strains after overnight incubation for 18 h was calculated relative to the WT strain 10403S. The ability of these strains to secrete GABA into the extracellular environment is shown (F). Statistically significant differences were determined by paired *t* test (two-tailed) (ns, not significant; * *p* < 0.05; **, *p* < 0.01; ***, *p* < 0.001).

### Adaptive Acid Resistance is highly GadR-dependent

Since GadR contributes significantly to acid resistance in stationary phase, we reasoned that it might also participate in regulating the ATR in *L. monocytogenes*. To test this, the transcription of *gadT2*, *gadR*, *gadD1*, and *gadD3* was measured in exponentially grown cultures of strains EGD-e wild type (WT) (*gadR*^-^) and the isogenic *gadR*^+^ derivative after 15 min exposure to a range of low pH conditions (pH 3-6.5, compared to pH 7). While *gadT2*, *gadD1*, and *gadD3* were all acid-inducible with a peak at pH 5, only the induction of *gadT2* was GadR-dependent (Fig 4A; S1 Fig). *gadR* transcription was unchanged regardless of the pH likely indicating that the *gadT2* induction might involve post translational regulation of GadR (Fig 4B). pH 5 activates *gadT2* transcription to the greatest extent and this activation appeared to be rapid and continuous (Fig 4C). The transcription of the SigB-dependent *gadD3* and *lmo0913* was also induced, however to a lesser extent (Fig 4C). These data demonstrated that GadR was responsible for the activation of *gadT2* transcription in response to mildly acidic stress.

**Fig 4.**
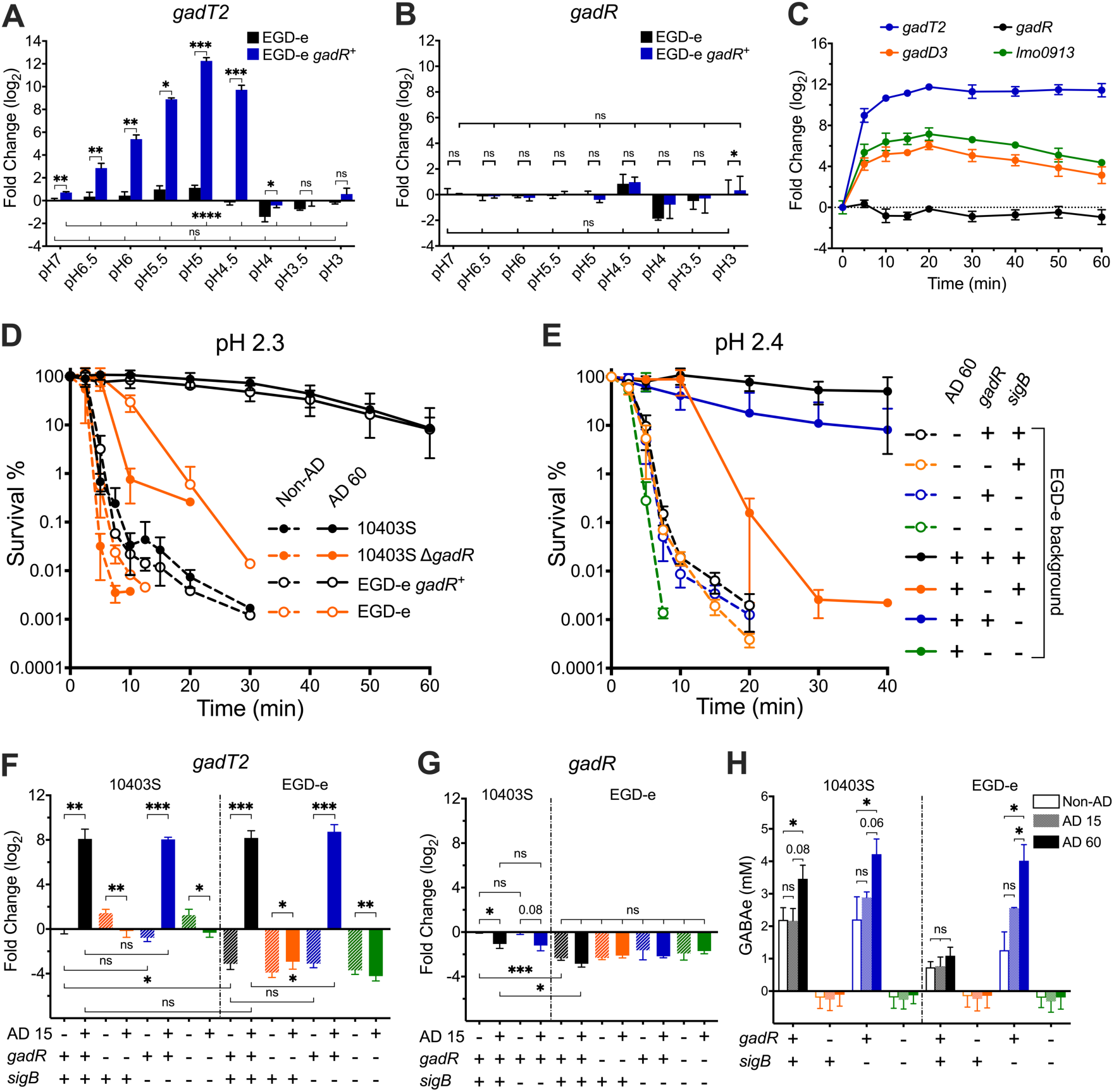
Adaptive acid resistance is GadR-dependent. The transcription levels of *gadT2* (A) and *gadR* (B) in exponential phase cultures of EGD-e WT (*gadR*^-^) or *gadR*^+^ strains with or without a 15 min exposure to pH 3.0 - 6.5 are shown, expressed relative to untreated EGD-e WT stain. (C) The differential transcription of *gadT2*, *gadR*, *gadD3*, and *lmo0913* in response to pH 5.0 exposure was monitored throughout a 60 min period in strain EGD-e *gadR*^+^. Cultures of *L. monocytogenes* WT or mutant strains grown to exponential phase with (AD 60) or without (Non-AD) a 60 min pH 5.0 adaptation were challenged in pH 2.3 (D) or pH 2.4 (E). Transcription of *gadT2* (F) and *gadR* (G) in exponential phase cultures of *L. monocytogenes* WT or mutant strains with or without 15 min exposure to pH 5.0 (AD 15) are calculated and shown relative to the untreated 10403S WT strain. (F) The GABA exported into the medium was measured using exponential phase cultures either untreated (Non-AD) or exposed to pH 5.0 for 15 min (AD 15) or 60 min (AD 60). Statistically significant differences between measurements were determined using paired *t* test (two-tailed) (ns, not significant; * *p* < 0.05; **, *p* < 0.01; ***, *p* < 0.001). Statistically significant differences across a group of samples were determined by one-way ANOVA (ns, not significant; * *p* < 0.05; **, *p* < 0.01; ***, *p* < 0.001).

To test the involvement of GadR in the ATR, the ability of strains 10403S and EGD-e to survive at pH 2.3 with or without GadR was measured before (Non-AD) and after pH 5 adaption for 60 min (AD 60). All strains from exponential phase without pH 5 adaptation exhibited low levels of acid resistance. Strains 10403S WT and EGD-e *gadR*^+^ developed a strong ATR after adaptation (Fig 4D and 4E), while strains 10403S Δ*gadR* and EGD-e WT (*gadR*^-^) acquired greatly reduced levels of acid resistance after pH 5 treatment (Fig 4D). Since EGD-e is the most widely studied strain, the relative contributions of *gadR* and *sigB* to ATR were examined using strain EGD-e WT (*gadR*^-^) and its isogenic mutants (*gadR*^+^, Δ*sigB*, *gadR*^+^ Δ*sigB*). Because of the extreme acid sensitive phenotype of strain *gadR*^-^ Δ*sigB*, the ability to survive at pH 2.4 (rather than pH 2.3) was measured to compare the ATR. The ATR was only slightly reduced following *sigB* deletion in strain EGD-e *gadR*^+^ but it was essentially absent when *sigB* was deleted in strain EGD-e (*gadR*^-^) (Fig 4E). In accordance with these observations, *gadT2* was strongly induced by pH 5 adaption for 15 min (AD 15) in both genetic backgrounds independently of SigB (Fig 4F). *gadR* transcription appeared to be slightly downregulated following adaptation in strain 10403S however it remained unchanged in EGD-e genetic background (Fig 4G), further suggesting that GadR activation by mild acidification was likely achieved through post translational regulation. Consistent with previous phenotypic and transcriptional observations, Glu/GABA antiporter activity was only present and induced in the *gadR*^+^ strains (Fig 4H). Interestingly, the ability to secrete GABA was notably higher in the absence of *sigB* in EGD-e strains (Fig 3F and 4H), although these differences were not reflected in the transcriptional or acid resistance measurements (Fig 3B and 5E). Taken together, these results suggested that exposure to pH 5 induces a strong GadR-dependent ATR in *L. monocytogenes*.

### Two 18 bp-palindromes in PgadT2 are essential for GadR-dependent ATR

Although *gadT2D2* alone confer GadR-mediated acid resistance, it remained unknown whether GadR controls the expression of genes that do not directly contribute to acid resistance. To test this, the global gene transcription during exponential growth with or without a 15 min pH 5 treatment was compared between strains EGD-e WT (*gadR*^-^) and *gadR*^+^. No gene was significantly differentially regulated without acid adaption between these two strains (S2 Fig). In contrast, 382 genes were upregulated and 374 genes were downregulated in response to pH 5 treatment in strain EGD-e *gadR*^+^ (Fig 5A and S2 Table), but only the acid induction of the *gadT2D2* operon and an adjacent gene *lmo2361* appeared to be dependent on GadR (Fig 5B). Moreover, *gadT2D2* were co-transcribed as a transcriptional unit and they were the most upregulated genes (11.7 and 9.3 log_2_ fold changes, respectively) in response to pH 5 treatment across the genome (Fig 5A and S2 Table), supporting a prominent role for GadR in the ATR. These data demonstrated that in response to mild acid stress GadR exclusively influences the transcription initiated at *lmo2361*-*gadT2* intergenic region.

**Fig 5.**
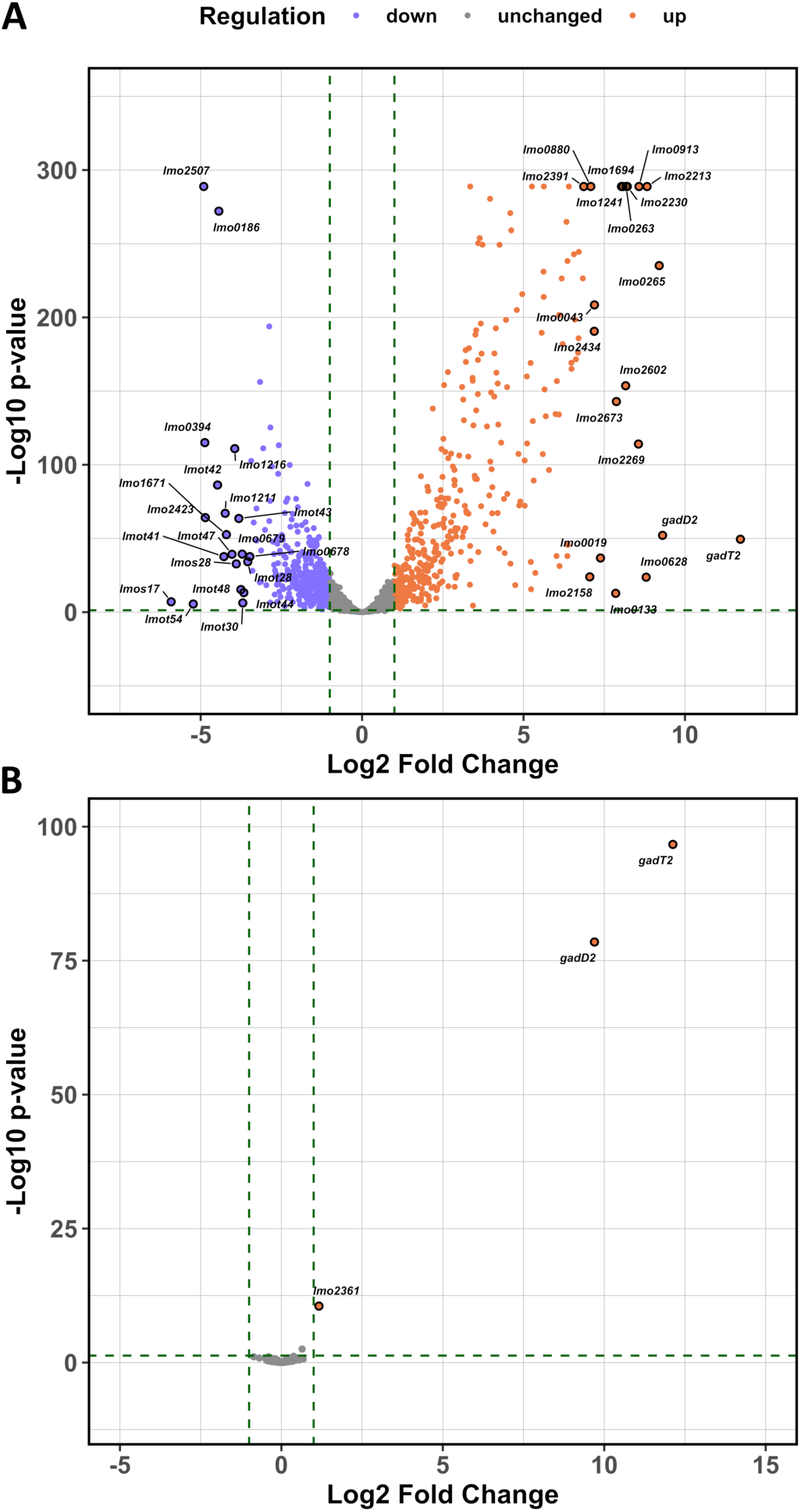
GadR is a dedicated regulator of the *gadT2D2* operon during acid adaptation. (A) Global gene transcription following a 15 min pH 5.0 treatment was measured using RNA-Seq, with values expressed relative to the untreated exponential phase culture of strain EGD-e *gadR*^+^. The 20 most upregulated/downregulated genes are labeled. (B) GadR-dependent genes were detected by comparing global gene transcription of strain EGD-e *gadR*^+^ to the WT EGD-e strain (*gadR*^-^) after 15 min at pH 5.0. Genes showing differential transcription are labeled. Genes with differential transcription > 2 fold and with *p*-value < 0.05 were considered significantly differentially regulated (marked by dotted lines on the Volcano plots).

To identify the sequence motif(s) required for GadR-mediated transcriptional regulation, P*gadT2* (defined as the sequence between the coding sequences of *gadT2* and the upstream adjacent gene) was carefully examined. Since the genes upstream from *gadT2* were not conserved across the *Listeria sensu stricto* clade (Fig 2), P*gadT2* sequence alignment was performed for *Listeria sensu stricto* spp. in order to search for conserved region, which is likely responsible for GadR-mediated *gadT2* regulation (Fig 6A). The transcription start site (TSS) of *gadT2* was mapped at ∼145 bp upstream of start codon using the RNA-seq data, a prediction that was also supported by previous results [40]. One notable feature that was conserved in all sequences was a pair of imperfect palindromic sequences that shared sequence similarity within the conserved region of P*gadT2* and upstream of TSS (Fig 6A and 6B). To test whether these two palindromes influence GadR-mediated *gadT2* transcription, a panel of mutants (designated V1 to V10) (Table 1) with various mutations in P*gadT2* was constructed (Fig 6A and 6B) and tested for *gadT2* transcription as well as acid resistance at exponential phase with or without pH 5 treatment. Deletion of the conserved region in P*gadT2* (V1, V2, and V3) abolished the GadR-mediated ATR (Fig 6). Disruption of either one of these two putative GadR-boxes (V4, V5, and V6) or the spacer region (V7) results in absence of GadR-mediated ATR (Fig 6). While a Δ1 bp deletion in the spacer between the two palindromic sequences (V8) was tolerable for GadR-mediated ATR, a 1 bp insertion in spacer region (V9) results in moderately reduced *gadT2* induction and absence of GadR-mediated adaptive acid resistance (Fig 6). The results also suggested that the sequence to the 5’ of these palindromes is responsible for *lmo2361* transcription (S3 Fig). Further examination of P*gadT2* revealed a putative peptide of 23 aa, proceeded by a consensus ribosome binding site (RBS) (GGAGG) (Fig 6A and 6B), although deletion of this RBS (V10) did not significantly affect the GadR-mediated ATR (Fig 6). Taken together, these data suggested that the pair of 18 bp-palindromes in P*gadT2* are likely GadR-binding boxes that are essential for the GadR-mediated ATR in *L. monocytogenes*.

**Fig 6.**
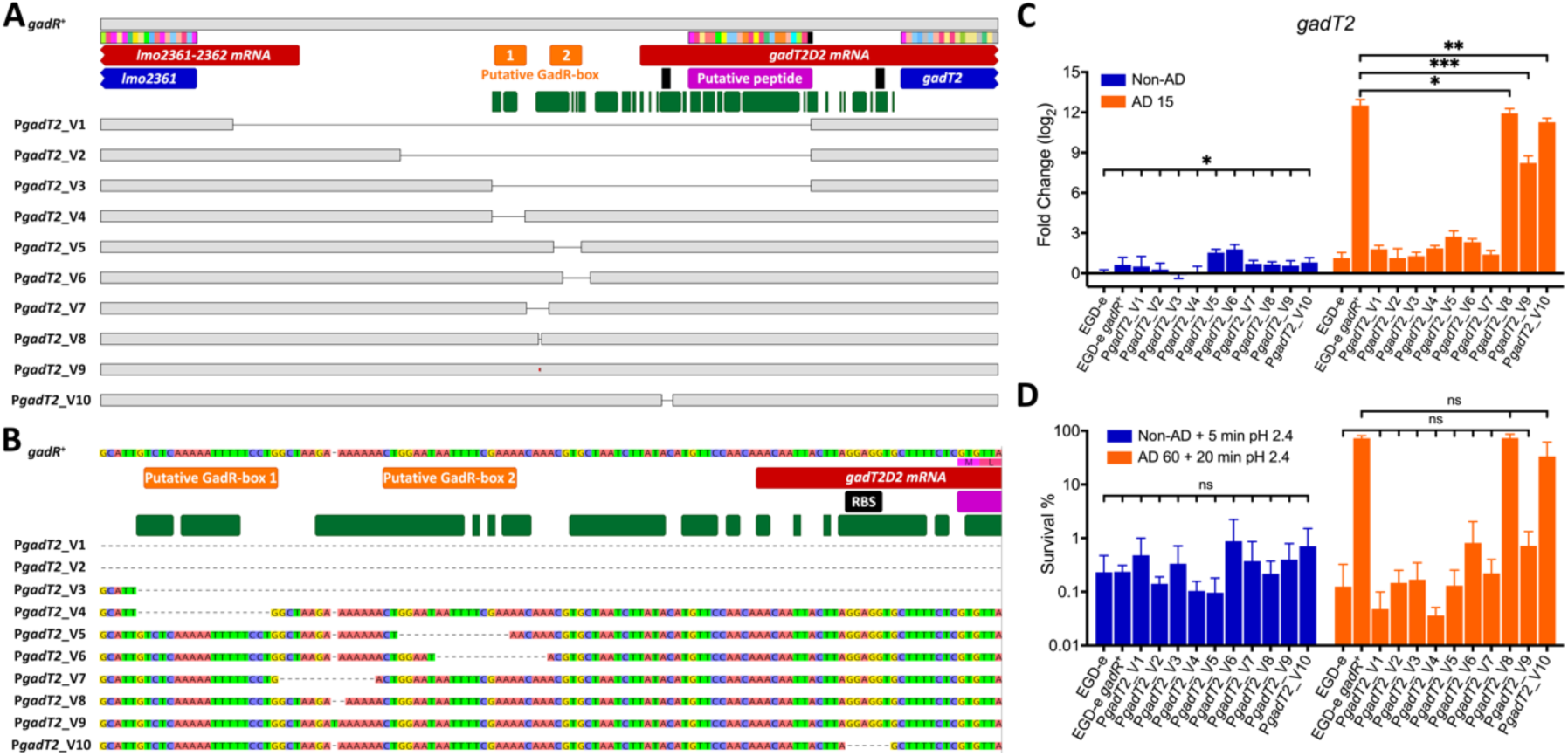
The promoter region of *gadT2 (*P*gadT2)* has two 18 bp-palindromes required for GadR-mediated activation of *gadT2D2* and induction of acid resistance. The intergenic sequence between the *lmo2361* and *gadT2* ORFs is depicted (A), with the nucleotide sequences shown for the 120 bp region targeted for mutagenesis (B). The sequences are annotated as follow: coding sequences are shown in blue; transcripts deduced from RNA-Seq analysis are shown in red; putative short translation frames are shown in pink; the putative GadR-boxes are shown in orange; predicted RBSs are shown in black; the 100% conserved bases across *Listeria sensu strico* spp. genomes that have *gadT2D2*-*gadR* gene cluster (n = 13, Fig. 1) are annotated with green boxes underneath. The mutants, labelled P*gadT2*_V1 to P*gadT2*_V10, were constructed in an EGD-e *gadR*^+^ background and their positions are indicated (A and B). (C) The transcription of *gadT2* was measured for the panel of P*gadT2* mutants during exponential phase either with (AD 15) or without (Non-AD) a 15 min pH 5.0 treatment and expressed relative to the untreated WT EGD-e strain (*gadR*^-^). (D) The ability of these strains to survive at pH 2.4 was determined using exponential phase cultures either with (AD 60) or without (Non-AD) a 60 min pH 5 treatment. Statistically significant differences between conditions were determined by paired *t* test (two-tailed) (ns, not significant; * *p* < 0.05; **, *p* < 0.01; ***, *p* < 0.001). Statistically significant differences across a group of samples were determined by one-way ANOVA (ns, not significant; * *p* < 0.05; **, *p* < 0.01; ***, *p* < 0.001).

## Discussion

In this study, a hitherto unknown RofA-like regulator, GadR was predicted by comparative genomics to contribute to acid resistance in *L. monocytogenes*. We have provided phenotypic, transcriptional, and biochemical evidence that GadR has a major impact on the adaptive acid resistance of *L. monocytogenes* by activating *gadT2D2* expression in response to a mild acid stimulus. Through transcriptomic analysis, we showed that GadR is a dedicated regulator of *gadT2D2* and identified two putative GadR-binding boxes that are essential for GadR-mediated acid resistance. The data presented here also help to explain the previously reported differences in GAD activity [23,28] and acid resistance [33,41] between strains of *L. monocytogenes*. The results suggest that adaptive acid resistance in this pathogen is largely controlled by GadR and SigB, the regulator of the general stress response. Overall, the study elucidates a key regulatory mechanism for adaptive acid resistance and helps to clarify the previously unresolved questions.

The conservation of *gadR-gadT2D2* in *Listeria sensu stricto* spp. (Fig. 1) points to the possibility that this acid resistance mechanism is important for the fitness of these species in the gastrointestinal tract, since they are commonly isolated from feces [42]. However, the occurrence of loss-of-functions mutation in *gadR* in strains EGD-e and 1381 raises the question of how common loss-of-function mutations in *gadR* are in *L. monocytogenes*. An inspection of the rate of PMSC occurrence in the *gadR* open reading frame per 100 bp in 40,080 sequenced *L. monocytogenes* genomes reveals that it is exceptionally high (S4 Fig). The PMSC rate in *gadR* is comparable to that of the *inlA* invasion gene, which has been reported to be highly susceptible to mutations resulting in truncation of the protein [43], and is an order of magnitude higher than the PMSC rates in *gadT2* and *gadD2* (S4 Fig). This finding suggests that allelic variation in *gadR* could be a major determinant of differences in acid resistance between *L. monocytogenes* strains. It is interesting to speculate about the evolutionary pressures that might lead to the selection of mutations in *gadR* despite its important in acid resistance. One possibility is that the GadT2D2 mediated reaction might be detrimental/dispensable under conditions where the availability of glutamate is limited or where maintenance of a high intracellular pool of glutamate is critical. Glutamate is present at a high concentration in the cytoplasm of *L. monocytogenes* where it contributes to osmoregulation and acts as a counter-ion for potassium [44]. It is conceivable therefore that under prolonged conditions of osmotic stress it might be advantageous to dispense with GadT2D2 by inactivating the main regulator. Alternatively, environments outside the mammalian host that rarely experience acidification might allow genetic drift of the *gadR* sequence, in the absence of positive selection for its function. Indeed, 8.4% of LII genomes encode truncated versions of GadR while this number is only 1.1% for LI genomes and even lower in the hypervirulent CCs: 0.3% in CC1, 1.1% in CC2, 0.2% in CC4 and CC6 [45]. Notably, a considerable percentage of strains from the phylogenetic groups that are prevalent in food are predicted to carry loss-of-function mutations in *gadR*, e.g. CC18 (86.3%), CC7 (31.4%), and CC9 (19.5%).

GadR shares significant sequence similarity with RofA, which was identified in *Streptococcus pyogenes* nearly three decades ago as a positive Regulator of protein F [46]. RofA is the founding member of a group of regulators called RALPs (RofA-like Proteins) that are found exclusively in Group A streptococci and that are a subset of the Mga superfamily of regulators [47,48]. Like the RALPs and Mga, GadR has two putative helix-turn-helix (HTH) DNA binding domains in the N-terminal half of the protein (Uniprot accessions HTH_Mga, PF08280 and Mga, PF05043). Both HTH domains of Mga are necessary for DNA binding and activation of the Mga regulon [49]. The presence of the two putative HTH domains in GadR suggests their involvement in binding to one or both of the predicted palindromic GadR-boxes upstream of *gadT2*. The predicted GadR-boxes span from -80 bp to -32 bp relative to the TSS of *gadT2* and GadR-box 2 likely overlaps with -35 box of the *gadT2* promoter, which is among the reported features of Mga superfamily transcription factors [50,51]. This finding suggests that when the GadR-box 2 is occupied by GadR the *gadT2D2* operon would be repressed whereas occupancy of the GadR-box 1, following an acid stimulus, would lead to derepression of the operon. Whether GadR dimerization occur remains to be established, although it is interesting to note that the GadR-mediated transcriptional activation of *gadT2* by acid is sensitive to the spacing between the two GadR-boxes; deletion of the spacer or insertion of 1 bp from it significantly reduces *gadT2* transcription following acid shock (Fig 6B and 6C) (V7 and V9, respectively). Based on our results, the likely scenario is that GadR is constitutively expressed regardless of the extracellular pH (Fig 4) and that some post-translational modification is induced by mild acid stress that facilitates/alters its interaction with the regulatory region. Two phosphotransferase regulatory domains (PRD) and a phosphotransferase system enzyme IIB-like domain are predicted at the C-terminal of GadR, similar to Mga and RofA, and these regulators were recently classified as PRD containing virulence regulators [52]. It is possible that the PRDs in GadR are involved in acid stress sensing and regulation of the activity of GadR. Further biochemical characterization of the GadR protein is underway to investigate this possibility.

While the GadR-mediated expression of GadT2D2 is shown to be a major factor in the development of an ATR, the global transcriptome analyses following pH 5.0 treatment reveals the complexity of the cellular response to acid. This acid stimulus was shown previously to trigger SigB activation [53] and indeed, ∼ 36% upregulated genes (n = 140) were SigB-dependent or proceeded by putative SigB promoter sequence (S2 Table). Moreover, the SigB regulon accounts for ∼ 86% (n = 60) of the 70 most upregulated genes (S2 Table), including genes involved in known acid stress response (e.g. arginine/agmatine deaminase) and virulence-associated mechanisms (bile resistance, glutathione biosynthesis and internalin expression) [20,54–58]. These data further demonstrate the importance of SigB in acid adaption. We have recently demonstrated that the trace metal (Mn^2+^ and Zn^2+^) homeostasis under acid stress is critical for an adequate stress response [34]. In accordance with this, the expression of Mn^2+^ importers (*lmo1424* and *lmo1847*- *1849*) and a putative Zn^2+^ exporter (*lmo2231*) was induced while the expression of putative Zn^2+^ importers was repressed (*lmo1445*-*1447* and *lmo1671*) (S2 Table). Several upregulated genes are involved in carbohydrate uptake and metabolism (S2 Table) indicating a general shift in carbon metabolism during growth in acidic conditions. Expression of multiple transcriptional regulators (e.g. *lmo2241*, *lmo2551*, *lmo0815*, and *lmo2494*) was also affected, and these might be partially responsible for the SigB independent differentially regulated genes (S2 Table).

The identification of GadR as the principal regulator of the GadT2D2 system and the characterization of global transcriptomic response to acid stress exposure help to understand the overall physiological response to mild acid stress in *L. monocytogenes*. Exposure to mild acid stress serves as signal to the rapidly growing cells to prepare for a harsher environment. The mostly prominent actions involve rapid GadR- and SigB mediated transcriptional response. GadR specifically promotes high expression of GadT2D2, which serves to protect against potentially lethal acid stress by helping to neutralize intracellular pH through glutamate decarboxylation. The total GAD activity is possibly supported by SigB-mediated *gadD3* expression, while other proton-consuming mechanisms (e.g., arginine/agmatine deaminase) might also contribute to the overall acid resistance [20]. SigB as the general stress response regulator promotes an array of mechanisms that contribute to intracellular pH maintenance (S2 Table) [19,30]. Our data suggest that there is an additive (rather than synergistic) effect in acid resistance between GadR and SigB. Besides intracellular pH maintenance, the cells also exquisitely manage the metal homeostasis and accelerate carbon source utilization under acid stress, presumably to avoid metal intoxication and to acquire energy. The acid exposure also serves as a signal for host entry for the bacterium to express apparatus that are essential for GI tract survival, intestinal epithelial adhesion and virulence factor activation [19] as supported by our transcriptomic data (S2 Table).

Overall, this study sheds new light on the regulatory mechanisms that underpin the acid stress response of this important food-borne pathogen. It further highlights the central role that glutamate decarboxylation plays in adaptive acid resistance. Further studies are underway to address some of the most important outstanding questions. Chief amongst these is the nature of signal detected by GadR in response to acidification. It could be either acid pH itself or some secondary effect of reduced pH on the physiology of the cell. It seems likely that some post-translational modification of GadR is required to activate it, but further work will be needed to clarify this. It will be interesting to learn whether GadR plays a role in colonizing the mammalian host, in particular if it contributes to surviving the transition through the acidic conditions in the stomach. If it proves to be critical for this early stage of the infectious cycle it might make a good diagnostic target for identifying strains of concern in food processing environments and potentially could provide a basis for novel strategies to control this pathogen.

## Materials and Methods

### Strains and culturing conditions

*L. monocytogenes* and *E. coli* TOP10 strains and plasmids used in this study were listed in Table 1. *L. monocytogenes* strains were grown in BHI (LAB M LAB048) at 37°C with agitation 150 rpm, unless otherwise specified. For stationary phase culture, an isolated *L. monocytogenes* colony was inoculated to 5 mL BHI broth in 50 mL centrifuge tubes and incubated for 18 h. To prepare exponential phase culture, overnight culture of *L. monocytogenes* was washed twice with fresh BHI broth and inoculated to 5 mL BHI broth in 50 mL centrifuge tubes to achieve initial OD^600nm^ = 0.05 and incubated for ∼ 3 h until mid-exponential phase (OD^600nm^ = 0.4). *E. coli* strains were grown in Luria-Bertani (Sigma). The following antibiotics were added to the medium where specified: kanamycin 75 μg · mL^-1^ (^kan^), ampicillin 100 μg · mL^-1^ (^amp^), chloramphenicol 10 μg · mL^-1^ (^chl^), and erythromycin 2 μg · mL^-1^ (^ery^).

### Molecular techniques

Plasmids and primers used in this study are listed in Table 1, 2 and S1. To complement *gadR* in strains 1381 and 1381R1, the full length *gadR* coding sequence was cloned (Phusion, ThermoFisher) from closely related CC2 strain 1380. The PCR product and expression vector pIMK3 [37] were digested (FastDigest, ThermoFisher) and ligated (T4 DNA ligase, Roche). 3 μL ligation mixture was used in the thermos-shock transformation of *E. coli* TOP10 chemically competent cells, the cells were plated on LB^kan^ plates after recovery and incubated overnight at 37°C. Transformant carrying correct insert in the plasmid was grown overnight in LB^kan^ to propagate the plasmid for purification. Both purified plasmid (pJW2) and empty vector (pIMK3) were electroporated into electrocompetent cells of *L. monocytogenes* strains 1381 and 1381R1 as previously described [19,37]. To construct EGD-e *gadR*^+^ and 10403S Δ*gadR*, two pMAD derivatives were constructed each with a synthesized (Eurofins) 600 bp insert containing 300 bp upstream and 300 bp downstream of mutation site with restriction sites introduced on both ends. For EGD-e *gadR*^+^, the *gadR* nonsense mutation in strain EGD-e was reverted in the insert (*374L) and several silent mutations were also introduced to facilitate the discrimination of mutant from WT using PCR [19]. The constructed plasmids were transferred to electrocompetent *L. monocytogenes* EGD-e or 10403S cells and spread onto BHI^ery^ plates as previously described [19,37]. The mutation was achieved by a two step integration [19,59]. The mutant and WT were discriminated by PCR. For introducing *sigB* deletion, the previously constructed pMAD Δ*sigB* was transfer to strain 10430S Δ*gadR* and EGD-e *gadR*^+^ [38]. To introduce Δ*gadT2D2R* and Δ*gadT2D2* in strain 10403S and to create mutant panel EGD-e *gadR*^+^ P*gadT2*_V1-V10, plasmids used for mutagenesis were constructed by SOE. Deletion of the entire P*gadT2* sequence was firstly introduced to EGD-e *gadR*^+^ to obtain EGD-e *gadR*^+^ P*gadT2*_V1. Various length of P*gadT2* were then complemented to EGD-e *gadR*^+^ P*gadT2*_V1 by two-step integration to obtain EGD-e *gadR*^+^ P*gadT2*_V2-V10.

### Transcriptional analysis

Transcriptional analysis was performed similar to previously described with minor adjustments [19]. Briefly, for stationary phase transcription analysis, RNA samples were taken directly from overnight culture. In acid adaption analysis, 5 M HCl was added to exponential phase culture (OD^600nm^ = 0.4) to acidify the media to pH 6.5, 6.0, 5.5, 5.0, 4.5, 4.0, 3.5, and 3.0 (the volumes of 5 M HCl required were pre-determined). RNA samples were taken for transcription analysis after 15 min of exposure to acid stress. Alternatively, the exponentially growing culture was acidified to pH 5 and incubated at 37°C. RNA samples were taken at 0, 5, 10, 15, 20, 30, 45, and 60 min for transcription analysis. To extraction RNA, 1 mL of culture was mixed with 5 mL RNALater (Sigma) and incubated at ambient temperature (∼18°C) for RNA preservation. All subsequent RNA extraction and cDNA synthesis steps were carried out at 4°C as previously described [33]. Cells were then recovered by centrifugation and resuspended in RLA buffer (Qiagen RNeasy minikit). The mixture was transferred to lysis matrix B tubes for mechanical lysis (40 s at 6 m · s^-1^, twice). The rest of RNA extraction procedures followed manufacturer’s instructions. The resulting RNA samples were treated with DNase (Turbo, Invitrogen). The quantality and quality of RNA samples were examined using Nanodrop before reverse transcription (SuperScript III, Invitrogen) was carried out (13 μL reaction, ∼ 0.5 μg RNA applied). qPCR (LightCycler® 480 SYBR Green I Master, Roche) was performed with 100 × diluted cDNA samples in a Roche LightCycler 480 system [33]. All primers for gene expression analysis were designed to anneal to regions that were conserved across strains to be analysed (Table 2). Three independent experiments were carried out each with two technical repeats. Relative gene transcriptions were calculated using Q-gene [60] with 16S as the reference gene.

**Table 2.**
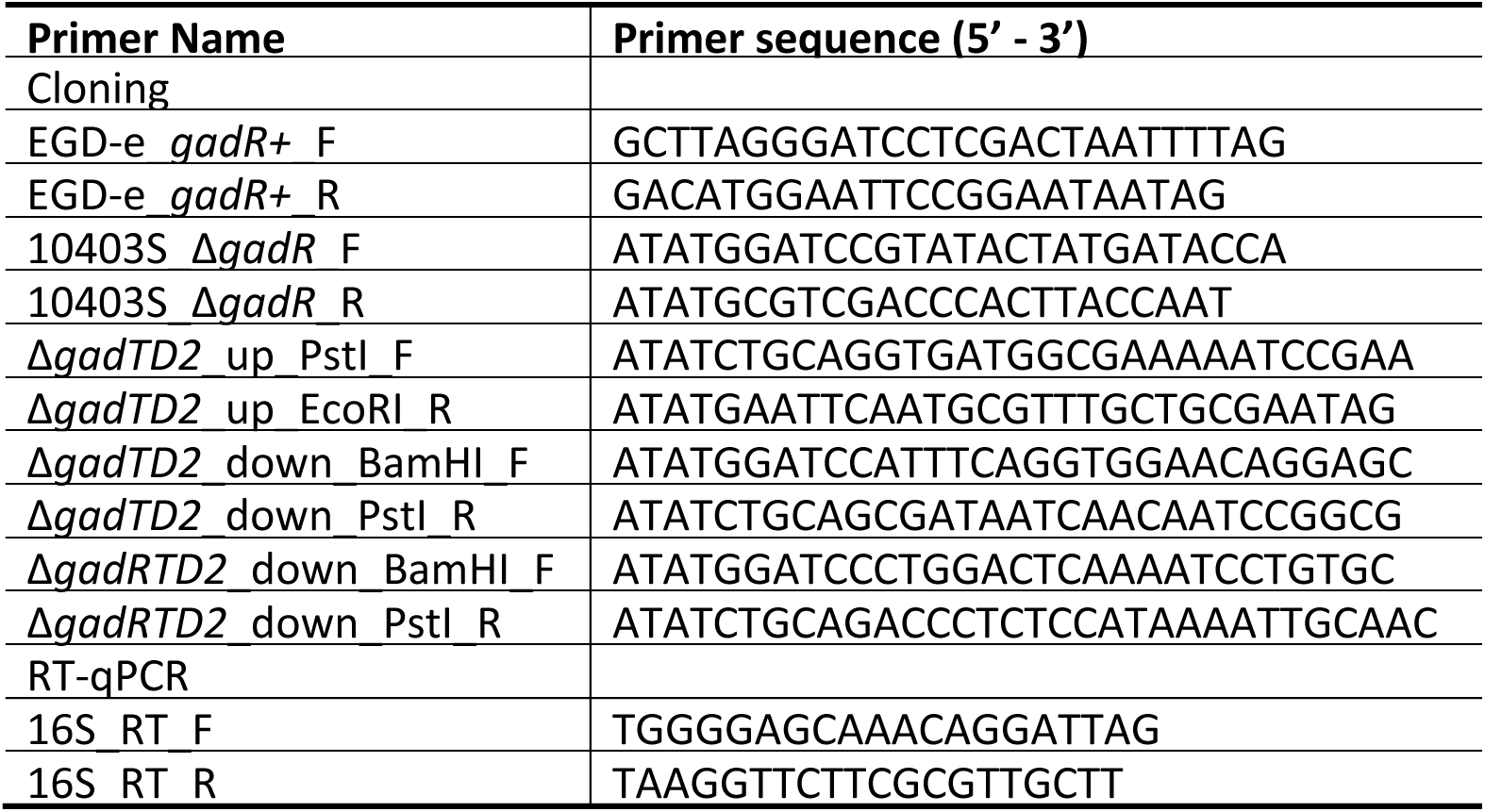
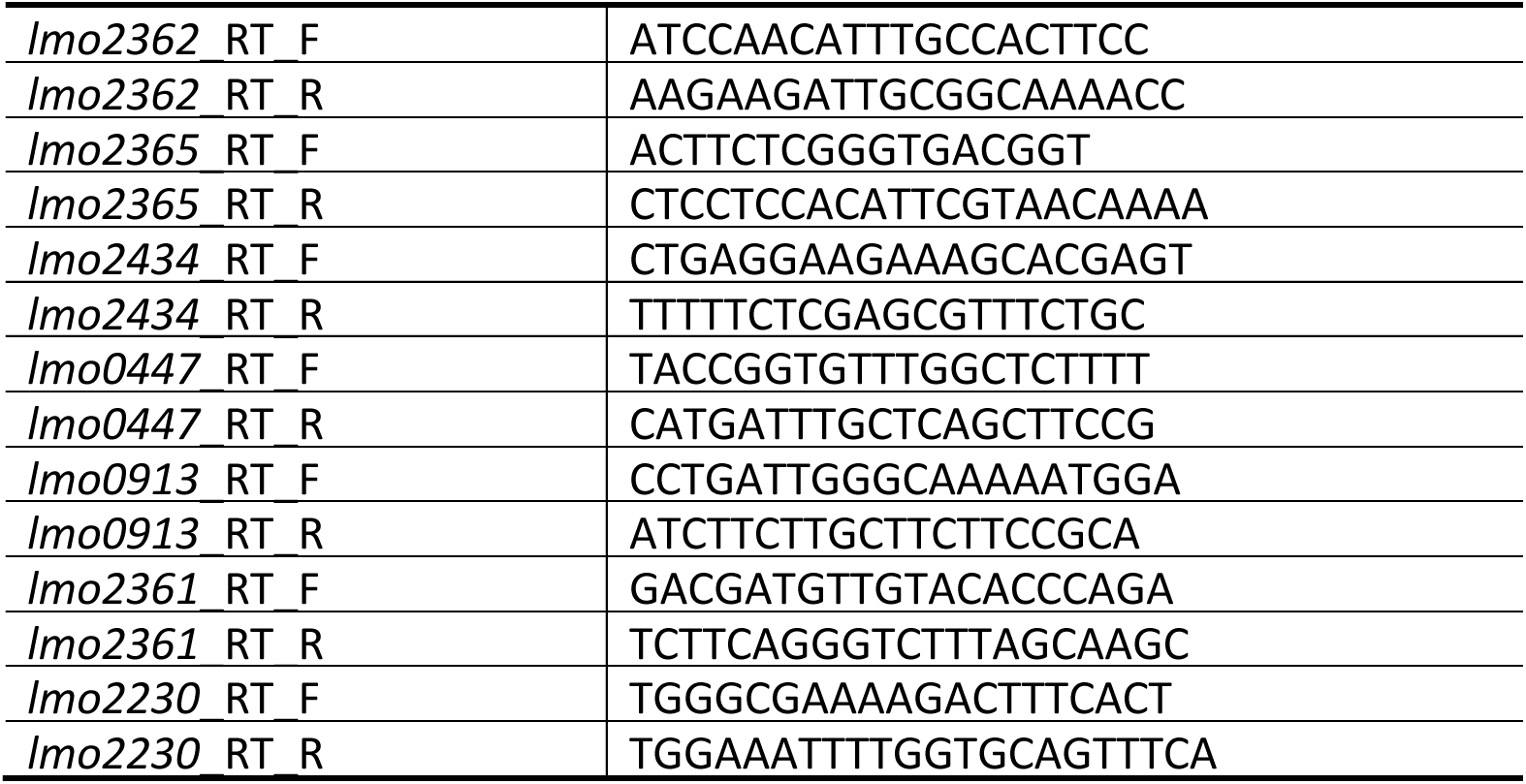
Primers used in this study.

### Acid survival experiments

The ability of *L. monocytogenes* strains to survive at lethal acidic condition was tested as previously described with minor adjustments [61]. To examine the acid resistance at stationary phase, 100 μL stationary phase culture was mixed with 900 μL BHI pH 2.3 or pH 2.4 in 1.5 mL microcentrifuge tubes and incubated at 37°C statically. Samples were taken at indicated time points, serial diluted (10 ×, in Phosphate-Buffered Saline) and 10 μL of cell suspensions from each dilution were then spotted onto BHI agar to quantify viable cells. Acid resistance was determined as the percentage survival at each time point relative to the starting cell count. Three independent experiments were carried out, each with triplicates. To examine the GadR mediated adaptive acid stress response, 100 μL exponential phase culture with/without acid adaption for 60 min was mixed with 900 μL BHI pH 2.3 or pH 2.4 in 1.5 mL microcentrifuge tubes. The samples were incubated and analysed following the same procedures as described for stationary phase acid resistance determination.

### GABA assay

The abilities of strains 10403S and EGD-e and their derivatives to produce extracellular GABA at stationary phase and exponential phase with/without acid adaption were examined using a previously established method with minor adjustments [62]. For stationary phase culture, 4 mL overnight culture was resuspended in 2 mL BHI^chl^ pH 3 and incubated for 1 h at 37°C. For adaptive acid stress response, 2 mL culture was taken from exponential phase cultures that were exposed under pH 5 stress for 0, 15, and 60 min then resuspended in 200 μL BHI^chl^ pH 3.25 and incubated for 1 h at 37°C. Following this incubation, the samples were span down (10,000 g × 5 min) and supernatant was collected for analysis. To quantify the GABA in the supernatant collected, 5 μL sample was added to 95 μL freshly prepared reaction master mix (80 mM Tris buffer, 750 mM sodium sulfate, 10 mM dithiothreitol, 1.4 mM NADP+, 2 mM α-ketoglutarate, and 0.1 g × L^−1^ GABase) and incubated at 37°C for 1 h in a temperature-controlled plate reader with OD340_nm_ measure every 60 s for 3 h. Standard curves were made by analysing 1-10 mM GABA solution (made with BHI pH 3 or pH 3.25). OD_340nm_ measured at 1 h was inspected for GABAe calculation. Three independent experiments were carried out, each with duplicates.

### Transcriptomic analysis

RNA samples for transcriptomic analysis were extracted as above mentioned and sent to Novogene for RNA sequencing and subsequent bioinformatic analysis following standard procedure. Birefly, for library construction ribosomal RNA was removed from total RNA with the Illumina Illumina Ribo-Zero Plus rRNA Depletion Kit. After fragmentation, the first strand cDNA was synthesized using random hexamer primers. During the second strand cDNA synthesis, dUTPs were replaced with dTTPs in the reaction buffer. The directional library was ready after end repair, A-tailing, adapter ligation, size selection, USER enzyme digestion, PCR amplification, and purification with AMPure XP beads. The library was checked with Qubit and real-time PCR for quantification and bioanalyzer for size distribution detection. Quantified libraries will be pooled and sequenced on Illumina platforms, according to effective library concentration and data amount required. The sequencing was performed in the Illumina NovaSeq6000 using a sequencing strategy based on paired end reads with a sequencing length of 150 bp per read (PE150). The raw reads were mapped to reference genome EGD-e, and log2 fold changes were calculated comparing the normalized readcounts between two given groups. Raw sequencing data is available from NCBI SRA under project number: PRJNA947476.

### Statistics

All statistical analysis were performed in Prism 8.

## Acknowledgments

The authors are grateful to colleagues in the Bacterial Stress Response Group at the University of Galway, Prof. Cormac Gahan (APC Microbiome, University College Cork, Ireland), Prof. Jörgen Johansson (Department of Molecular Biology, Umeå University, Sweden), and Dr. Lilliana Radoshevich (Department of Microbiology and Immunology, University of Iowa, United States) for helpful discussions. This project was supported by the Irish Department of Agriculture, Food and the Marine (17/F/244) and by the Science Foundation Ireland Frontiers for the Future Programme (21/FFP-P/10078). Jialun Wu was also supported by Irish Higher Education Authority funded Cost Extensions for Research Disrupted by COVID-19.

## Supporting information

**Fig S1.** *gadD1* and *gadD3* are both induced by acid stress independently of GadR.

**Fig S2.** Acid stress results in global transcriptomic response in strain EGD-e while GadR does not play a significant role in gene transcription under exponential phase.

**Fig S3.** *lmo2361*::*gadT2* intergenic sequence to 5’- end of putative GadR-boxes are required for *lmo2361* transcription.

**Fig S4.** High frequency of PMSCs was detected in the *gadR* coding sequence.

**Table S1** Plasmids and Primers used for creating mutants *L. monoctyogenes* EGD-e *gadR+* P*gadT2*_V1-V10.

**Table S2** Differentially expressed genes in strain EGD-e *gadR*^+^ vs wild type after acid treatment.

## Notes

### Competing Interest Statement

The authors have declared no competing interest.

